# Assessing the Role of Calmodulin’s Linker Flexibility in Target Binding

**DOI:** 10.1101/2021.03.15.435522

**Authors:** Bin Sun, Peter M. Kekenes-Huskey

**Affiliations:** Department of Cell and Molecular Physiology, Loyola University Chicago, Maywood, IL, USA 60153

## Abstract

Calmodulin (CaM) is a universal Ca^2+^ binding protein known to bind at least 300 targets. The selectivity and specificity towards these targets are partially attributed to the protein’s flexible alpha-helical linker that connects its N- and C-domains. How this flexible linker mediates the driving forces guiding CaM’s binding to regulatory targets is not well-established. Therefore, we utilized the Martini coarse-grained (CG) molecular dynamics simulations to probe interrelationships between CaM/target assembly and the role of its linker region. As a model system, we simulated the binding of CaM to the CaM binding region (CaMBR) of calcineurin (CaN). The simulations were conducted assuming a ‘wild-type’ calmodulin with normal flexibility of its linker, as well as a labile, highly flexible linker variant. For the wild-type model, 98% of the 600 simulations across three ionic strengths adopted a bound complex within 2 *µ*s of simulation time; of these, 1.7% sampled the fully-bound state observed in experimentally-determined crystallographic structure. By calculating the mean-first-passage-time for these simulations, we estimated the association rate to be *k*_*a*_ = 5.9 × 10^8^ M^*−*1^ s^*−*1^, which is similar to the experimentally-determined rate of 2.2 × 10^8^ M^*−*1^ s^*−*1^ [1]. Further, our simulations recapitulated the inverse relationship between the association rate and solution ionic strength reported in the literature. In contrast, although over 97% of the labile linker simulations formed tightly-bound complexes, only 0.3% achieved the fully-bound configuration. This effect appears to stem from a difference in the ensembles of extended and collapsed states controlled by the linker properties. Specifically, the labile linker variant samples fewer extended states compatible with target peptide binding. Therefore, our simulations suggest that variations in the CaM linker’s propensity for alpha-helical secondary structure can modulate the kinetics of target binding. This finding is important, as the linker region houses several CaM variants sites for post-translational modifications, that may alter the protein’s normal regulatory functions.

## 2 Introduction

Calmodulin (CaM) is a highly-expressed, 16.7 kDa globular protein comprising two domains connected by a linker (Fig. 1). CaM regulates hundreds of protein targets [2] in a Ca^2+^-dependent manner. How CaM maintains selectivity towards these targets with varying affinities (*K*_*d*_ from nM [3] to *µ*M [4]) has been studied for decades [5–16]. These studies have generated valuable insights into factors contributing to its binding selectivity, which includes hydrophobic anchoring residues from targets that interact with CaM, CaM’s conformationl heterogeneity at the binding surface and its Ca^2+^-binding sensitivity. Of these, the flexibility of its linker is believed to play a prominent role in shaping the conformational ensemble it adopts [17, 18] as well as its regulation of target enzymes [19].

**Figure 1:**
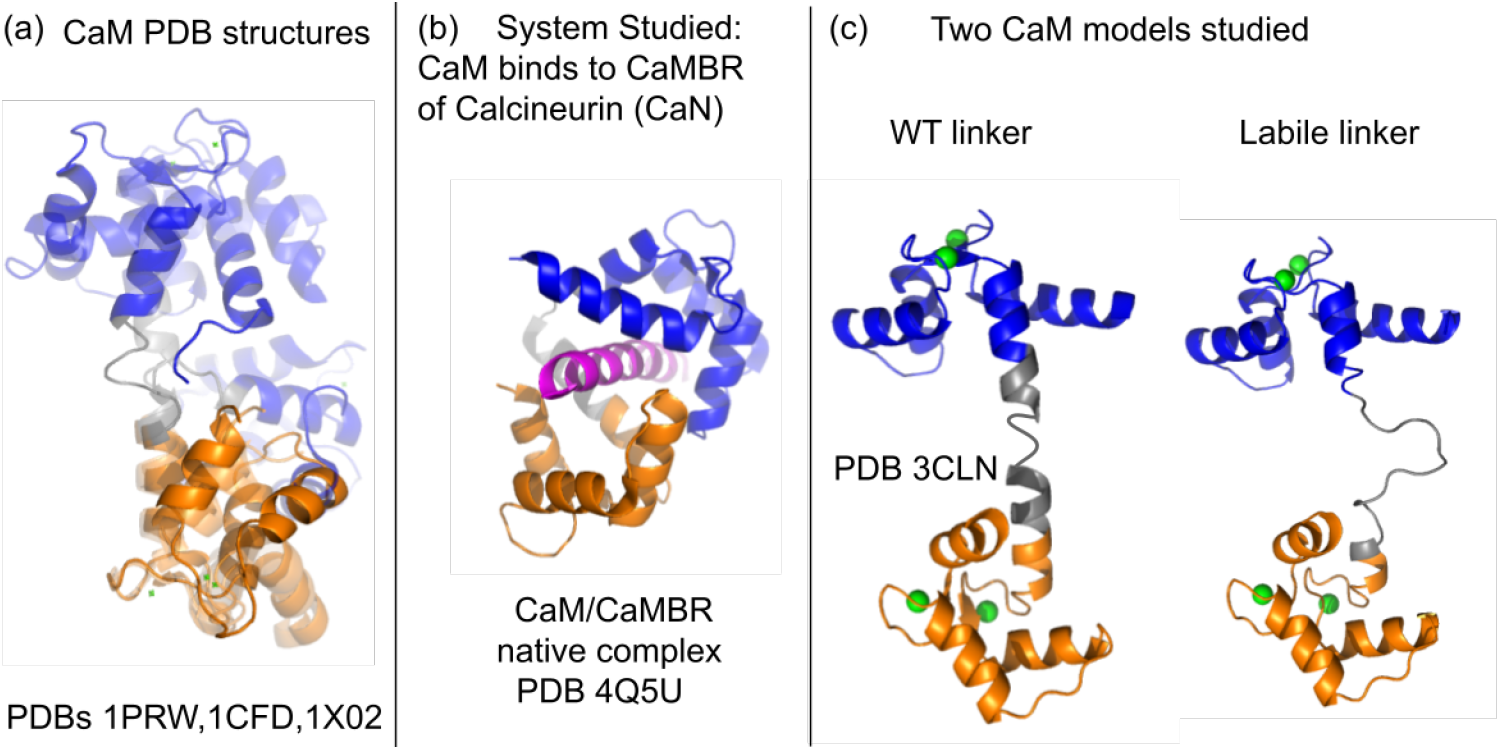
a) Three CaM crystal structures aligned on C-domain to show the native flexibility of the linker [2]. The N- and C-domains are colored blue and orange, respectively. The central linker (residues 73 to 87) is colored gray. The PDBs of these structures are 1PRW, 1CFD and 1X02. (b) The process of CaM binding to the CaM binding region (CaMBR) of calcineurin (CaN) is studied in this work. (c) The two CaM models that has a WT linker and a labile, highly-flexibilble linker, respectively, were studied in present work..

A key basis for CaM’s selectivity is attributed to the variety of binding modes it adopts when bound to its mostly peptide-based targets (reviewed in [20]). These include an extended conformation, a collapsed conformation in which its N- and C-domains wrap around the target, or intermediate configurations that permit CaM/target stoichiometries of 1:1, 2:2, or 2:1. Therefore, it is apparent that CaM’s ability to bind various targets in part stems from the linker’s ability to assume different conformations [21, 22]. Further, this flexible linker is believed to exert an entropic role in tuning target affinity, as target-binding induced mobility changes of linker residues correlate with the conformational entropy measured for the entire protein [22]. As an example, Katyal et al [23] demonstrated that tethering the N- and C-CaM termini via disulfide bond increases the entropic favorability of binding by reducing the ensemble of thermodynamically-accessible states available to the linker.

Nonetheless, while many thermodynamic details of CaM/target binding are increasingly understood, the protein’s binding kinetics remain enigmatic. Toward addressing this gap, molecular dynamics (MD) simulations have commonly been used to probe the CaM/target binding process. One of the prominent challenges in these simulations is the fact that most CaM targets are intrinsically disordered peptides (IDP) [24]. Such IDPs adopt stable secondary structures upon binding through “coupled binding and folding” mechanisms that are case-dependent [25] and generally a combination of conformational-selection and induced-fit [26, 27] binding strategies. IDP binding mechanisms can unravel over micro-seconds and longer time scales that are generally inaccessible to conventional all-atom protein descriptions [28]. Efforts to bridge this limitation include adaptive sampling techniques such as goal-oriented sampling, weighted ensemble (WE) and coarse-grained techniques, which have been used to study the process of IDP/DNA binding [29], globular proteins binding [30] and protein/ligand binding [31]. Of these approaches, coarse-grained (CG) molecular dynamics simulations are attractive due to their sampling efficiency and ability to preserve important molecular details of IDP-binding [32–36]. Such simulations could provide important insights into the time-dependent nature of CaM/target assembly, but have generally been limited to study their mechanisms, not kinetics [12, 33]. Resolving calmodulin (CaM) binding kinetics via coarse-grained modeling could shed insight into fundamental properties that determine CaM’s ability to regulate targets. This strategy could be important in resolving how known missense mutations [37] and post-translation modifications [38] in CaM’s linker domain may influence its function.

In this study, we simulated the binding process of a model system, CaM and the CaM binding region (CaMBR) of calcineurin, using a Martini CG model with explicit water molecules and ions. This permitted us to sample native-like CaM/CaMBR complex structures from unbiased sampling over a range of ionic strengths and under different linker flexibilities. Using these simulations, we have identified a mechanistic basis for how CaM linker flexibility shapes the kinetics of target binding and its dependence on the solvent ionic strength.

## 3 Materials and methods

### 3.1 Martini CG simulations

We elected to use Martini CG given its advantages over the ‘structure based’ model (SBM, also called G*o*-like model [39]). 1) Martini CG has higher molecular resolution than G*o*-based CG model. The Martini CG mapping ratio is on average four atoms to one CG bead [40] while the commonly used G*o*-based model are about ten non-hydrogen atoms to one bead [41]. This higher resolution permits detailed description of protein side-chain interactions with waters, which can impact protein mobility [42]. 2) No reference-structure based potential is needed, therefore the Martini CG potential have better transferability to arbitrary proteins without refitting and the simulation time scale can be directly interpreted. Therefore, Martini CG is well-suited for modeling the kinetics of CaM/target binding.

To this end, we established a Martini CG model based on the extended CaM structure PDB 3CLN [43]. The CaMBR of CaN is extracted from the CaM/-CaMBR complex PDB 4Q5U [44]. Both CaM and CaMBR structures were used to construct the CG model with the Martini protein force field V2.2 [45, 46]. Martini CG’s time scale is four times faster than all-atom simulations [47]. The mapping ratio we used assumed four heavy atoms to one bead on average. Martini CG does not typically sample protein secondary structure thus it relies on user-provided secondary structure information to assign proper backbone parameters of bonds, angle, and dihedral terms [45]. Therefore, we calculated the secondary structure of the CaM and CaMBR via the DSSP program [48, 49]. We constructed two CaM models, one has the WT linker (residues 73-87) flexibility and one has a labile, highly flexible linker (Fig. 2). The WT linker is based on PDB 3CLN and has a 4-residue hinge (residues 78-81) with DSSP predicted secondary structure as turns (“TTTT”). The labile linker in which the entire linker was subject to a short annealing MD in vacuum by Amber [50] to destroy the alpha-helix structure with DSSP predicted secondary structure as coils and bend (“CCCSCCSSSCCCSSS” where “C” and “S” refers to coil and bend, respectively). After defining the system, the CaMBR was randomly placed around CaM via the gmx insert-molecues command with a minimum distance between CaMBR and CaM of 40 Å to minimize bias.

**Figure 2:**
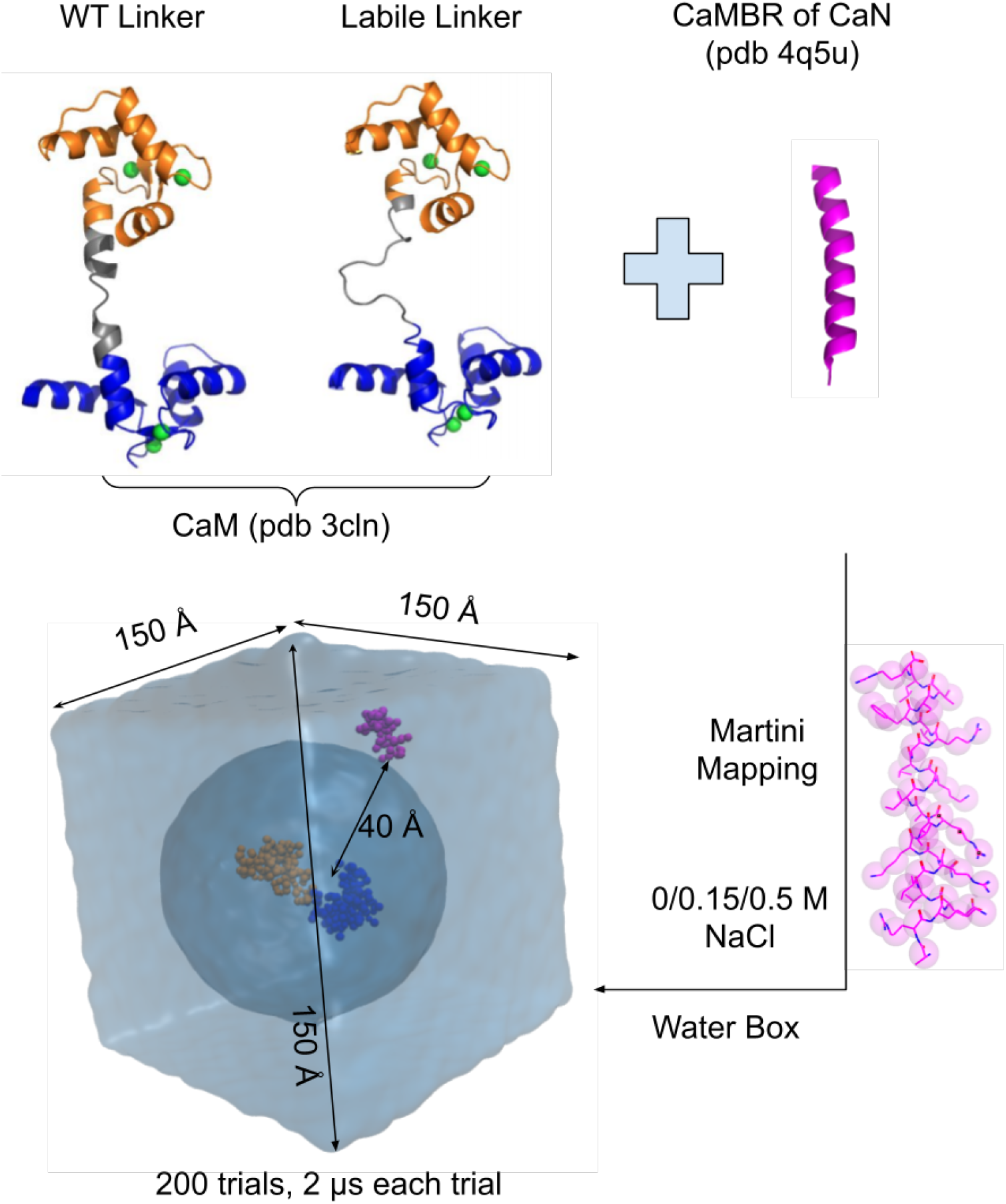
Martini CG setup. The all-atom CaM and CaMBR of CaN structures are from PDB 3CLN and PDB 4Q5U, respectively. Two CaM models with WT linker and labile linker were considered. After mapping the all-atom structures into Martini CG structures, the CaMBR was randomly placed around CaM with minimum distance *>*40 Å. The binding process was simulated by molecular dynamics without any further constraints imposed on the proteins..

For each system, 200 trials were run with ionic strengths of 0, 0.15, and 0.5M NaCl, respectively. Elastic constraints within CaM’s N-/C-domains were introduced to maintain an open domain conformation [51], which was justified by the observations that the N-/C-domain conformations are unchanged after binding CaMBR (*<* 2 Å RMSD). The Martini V2.0 ion force field was used to describe the NaCl ions added into the system and appropriate numbers were added to set the aforementioned ionic strengths. The system was solvated into a cubic water box with dimension of 150 Å per side using the gmx solvate command with pre-equilibrated waters from the Martini website. The standard Martini water model was used, in which four all-atom water molecules are combined into a single coarse-grained bead [52]. The system was first subject to 2000 steps of energy minimization using the steepest descent algorithm with constraints on the protein. The minimized system was equilibrated after being heated to 300 K over 6 ns. Constraints were imposed on the proteins during the heating stage. A 2 *µ*s production run was initiated from the equilibrated system in the NPT ensemble using a 30 fs time step. The Berendsen temperature and pressure couplings were used to maintain 300 K and standard pressure. All simulations were performed using GROMACS version 2020.3 [53]. The back mapping from CG model to the all-atom model was done according to the procedure proposed in [54].

### 3.2 Analyses

The autoimage command from the CPPTRJ [55] program that centers and images the trajectory was used for periodic boundary condition treatment. The trajectory was then converted to PDB format with bonds added using the gmx trjconv command of Gromacs. The PDB format trajectory was used for all analyses and visualization. The contacts between CaMBR and CaM were calculated assuming that one contact represents as any pair of beads that is within 5.5 Å. The CaMBR/CaM center of mass distance, RMSD to the native-like complex and CaM’s radius of gyration (*R*_*G*_) were calculated via the CPPTRAJ program. The CG structure of PDB 4Q5U served as a reference structure for the RMSD calculation. The trajectories were projected onto a plane according to the CaM/CaMBR center of mass distance and RMSD relative to the reference structure. The projection densities were estimated by a Gaussian kernel using the gaussian kde from the scipy python library. The potential was estimated via Boltzmann inversion −*k*_*b*_*T* ln(*ρ/ρ*_*min*_) where *k*_*b*_ is Boltzmann’s constant, *T* is temperature, *ρ* is the point density after projection and *ρ*_*min*_ is the minimum density.

### 3.3 Association rate calculations based on first passage time to bound state

CaM and CaMBR were deemed bound when the structures assumed 50 or more contacts. We later justify that CaMBR/CaM complexes by this definition are thermodynamically favourable and located near the fully-bound state (Fig. 5a). The first passage time (*T*_*fp*_) of reaching the bound state can then be used to estimate the association rate (*k*_*a*_) [56]. To illustrate the concept, we use the following expression:

**Figure 3:**
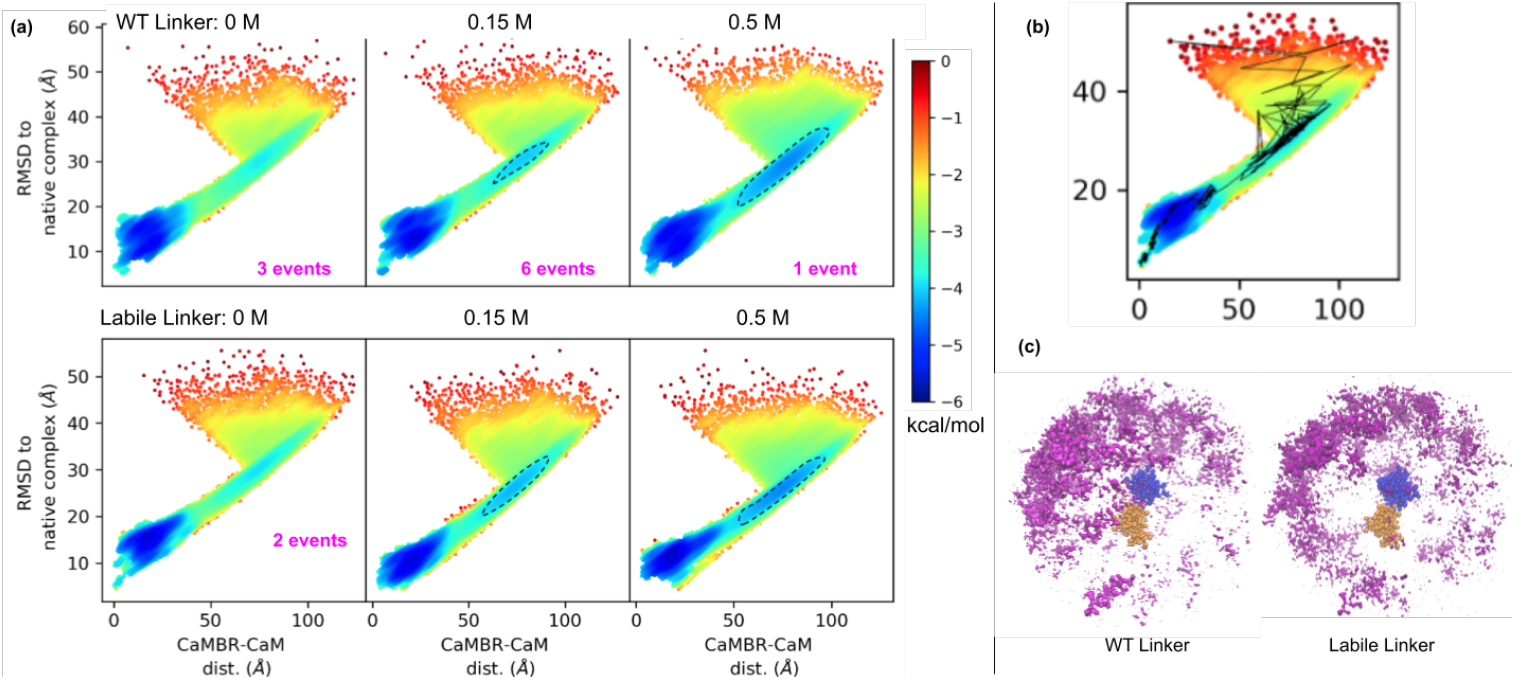
(a) Simulation trajectories projected onto 2D plane with axis: RMSD to native complex PDB 4Q5U and CaMBR/CaM center of mass distance. The dashed lines highlight the difference caused by increasing ionic strength. Energy is calculated via Boltzmann weights. The numbers of binding events that lead to native-like bound state are indicated in each panel. (b) An example trajectory of a binding event that leads to a native-like bound state. (c) Volumetric occupancy density isosurface (isovalue = 0.00015) of CaMBR beads around the CaM calculated using structures at the bottle-neck region indicated by dashed circles in panel a (RMSD∼ 30 Å and CaMBR/CaM distance ∼75 Å)

**Figure 4:**
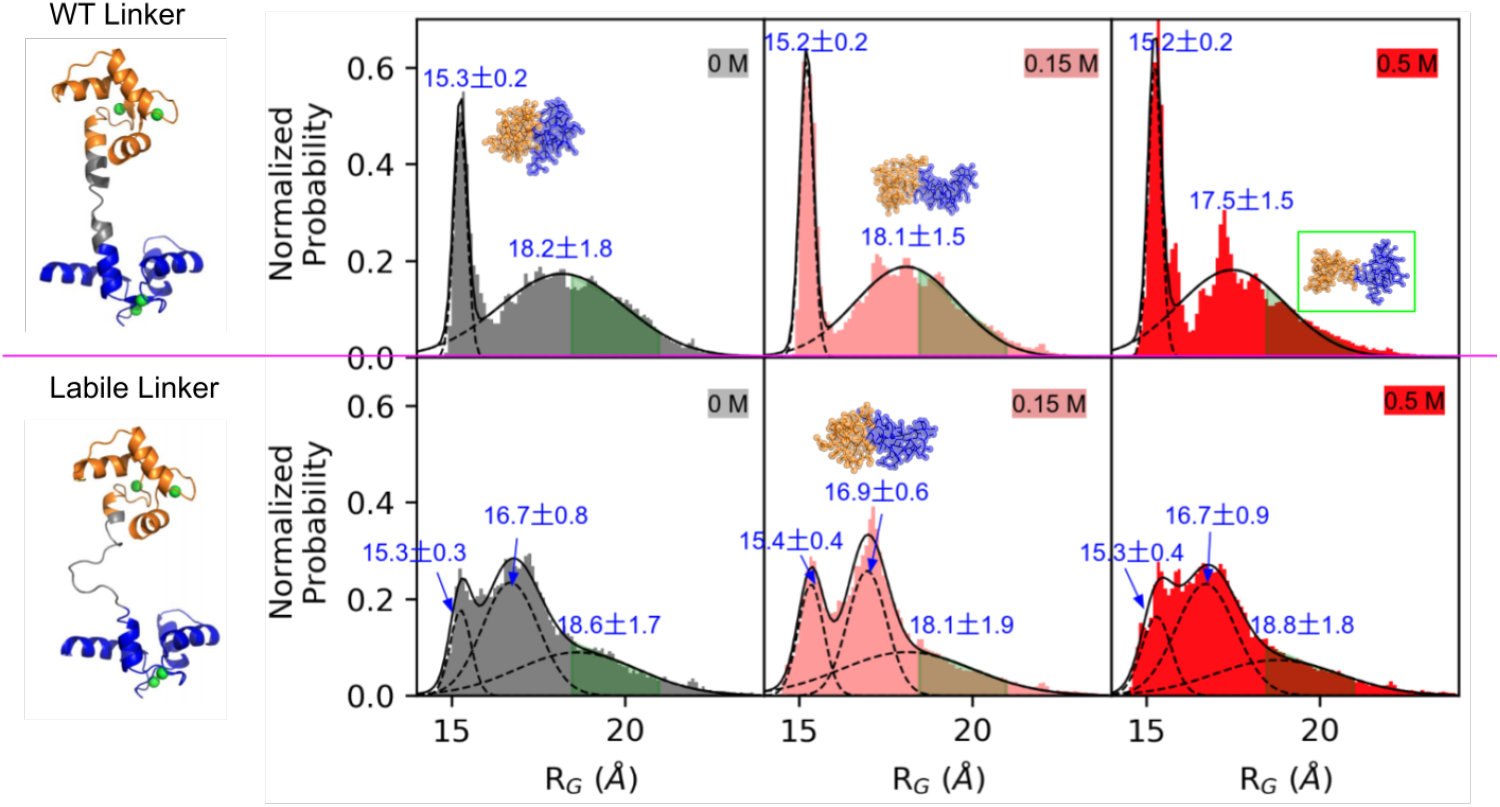
Radius of Gyration (*R*_*G*_) of CaM at different ionic strengths before forming the loosely-bound state with CaMBR. The probability is normalized such that the area under the curve is 1. The green shaded areas show the *R*_*G*_s of all 12 binding events that lead to native-like CaMBR/CaM complex with *R*_*G*_ = 19.7 ±1.3 Å. The three structures in the top row have *R*_*G*_ values of 15.2, 18.0 and 19.7 Å, respectively. The one structure in the bottom row has *R*_*G*_value of 17.0 Å..

**Figure 5:**
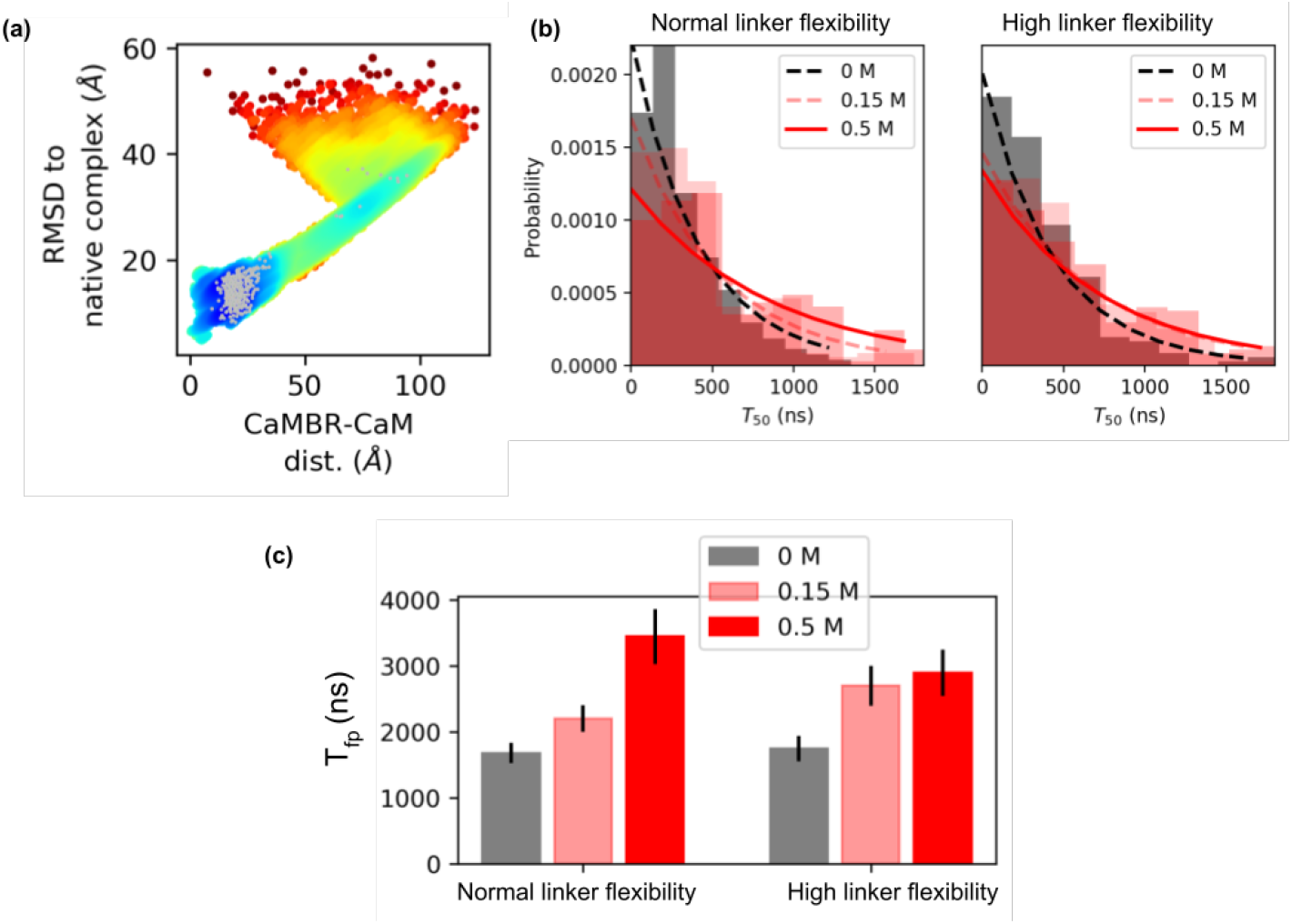
a) Locations of loosely-bound state complexes defined as 50 inter-contacts between CaMBR and CaM. b) Distribution of *T*_*fp*_s for a sample consisting of 200 simulation trials (with replacement). The distribution was fitted to *A* exp (*− τx*) and the expected *T*_*fp*_ is 1*/τ* ns. c) Average *T*_*fp*_ after bootstrapping using 1000 samples..

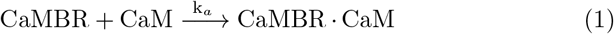

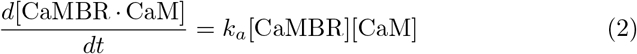

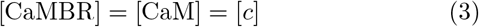

[*c*] = 1*/*(*N*_*A*_*V*_*box*_) is the equivalent concentration of one molecule in the simulation box with volume *V*_*box*_ = 3.375 × 10^*−*21^ L and *N*_*A*_ is Avogadro’s constant. In unit time (*dt* = 1), there are 1*/T*_*fp*_ association events, for which each event leads to the formation of CaMBR.CaM. Thus *d*[CaMBR · CaM] = (1*/T*_*fp*_)[*c*] = *k*_*a*_[*c*][*c*] which leads to:

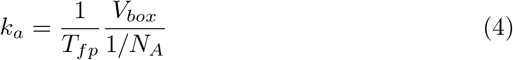

Hence, *k*_*a*_ is essentially the inverse of *T*_*fp*_ [56]. We used bootstrapping with replacement [57] to calculate the *T*_*fp*_ from our simulations. For this, we randomly generated 1000 distributions of *T*_*fp*_ values using the original 200 values generated by the simulations. Each of those 1000 distributions were fitted to an exponential distribution *A* exp (−*τx*), which is analogous to a Poisson process undergoing rare (binding) events. The expected value for *T*_*fp*_, **E** = 1*/τ*, is reported. Scripts for these analyses are available in https://github.com/huskeypm/pkh-lab-analyses (2021-CaMBRmartini)

## 4 Results and Discussion

### 4.1 Linker flexibility impacts CaM/CaMBR assembly

To investigate how the CaM’s linker flexibility affects the binding process between CaM and the CaMBR of CaN, we performed extensive binding simulations using the Martini CG models with two CaM models that have WT and labile linker flexibility, respectively. The control of linker flexibility is achieved by setting the secondary structural properties of the linker. Hence a linker with a higher helical character is more rigid than a coil. For the target-free CaM crystal structure (PDB 3CLN), the linker (residues 73-87) was anticipated to have moderate flexibility. This was based on the DSSP result [48, 49] suggesting that a four-residue ‘hinge’ of consisting of residues 78-81 assumed a ‘turn’-like (T) secondary structure, whereas the remainder were ascribed rigid alpha-helical character. For the labile linker we assumed the entire linker consisted of labile coils and turns. To study the contributions of long-range electrostatic interactions in driving diffusion [58, 59], we performed 200 simulation trials at 0, 0.15, and 0.5 M ionic strength, respectively. The trajectories were projected along two axis: a) the CaMBR/CaM center of mass distance b) the RMSD to native CaM/CaMBR complex crystal structure PDB 4Q5U (Fig. 3) to visualize the sampled conformational space. The population densities in the projected space was used to infer the potential of mean forceby inverting the Boltzmannequation, e.g., *E*_*PMF*_=−*k*_*b*_*T* ln(*ρ/ρ*_*min*_), where *k*_*b*_ is Boltzmann’s constant, *T* is temperature, *ρ* is the point density after projection and *ρ*_*min*_ is the minimum density.

Our simulations indicate that each system configuration exhibited significant sampling of a loosely-bound state, which is evidenced by the region of more negative *E*_*P MF*_ values found where the RMSD*<*20 Å and CaMBR/CaM distance*<* 35 Å. We note that the loosely-bound states most prominently sample a region modestly displaced from the reference crystal structure. We also observed some binding events that lead to the native-like bound complex. The resulting structures have an average RMSD of 6.3 Å to the crystal structure of the CaMBR/CaM complex; although this value is perceived as relatively large for atomistic-resolution structures, by visual inspection (Fig. S2) it is apparent that the trial poses strongly resemble the native structure. Additionally, the ∼6 kcal/mol potential difference of this region relative to the unbound state indicates that the binding process is thermodynamically favourable. Interestingly, a bottle-neck is also apparent when the RMSD is within 25 − 40Å and the CaMBR/CaM distance is between 60 − 100 Å for non-zero ionic strengths (see the dashed circles in Fig. 3a). We attribute this bottle-neck region to two factors: 1) Electrostatic screening by ions and 2) impaired CaM/CN alignment. We show in Fig. S3 that the electrostatic potentials about CaM and CaMBR are prominent, complementary and nonuniform. The largely negative electrostatic potential presented by CaM can be expected to facilitate the binding of the positively-charged CaMBR peptide, as is well-established in other protein/protein complexes [58]. However, the negatively-charged potential can also drive off-target associations of the two proteins that could compete with binding to the native bound configuration. With increasing ionic strength, it appears that the binding to the native bound configuration is disfavored, which manifests in a greater population of CaMBR/CaM binding poses that are off-target.

The nonuniformity of the CaM electrostatic potential also appears to impose constraints on the CaMBR’s ‘angle of approach’ (Fig. 3c). This is shown by the asymmetry in the distribution of CaMBR configurations about CaM. In other words, the nonuniform electrostatic potential of CaM and its highly labile conformational ensemble necessitates proper alignment of CaM with the CaMBR to facilitate productive binding. This mechanism of constraining the angle of approach has previously been observed in lysozyme/*α*-lactalbumin assembly [60]. In that study, it was shown the binding process for two charged protein substrates has a strong preference for a narrow set of approach angles [60, 61]. Further, the tendency for the two substrates to align decreased with increasing ionic strength. Therefore, we anticipate that the interplay of electrostatics and protein/protein alignment for CaM/CaMBR gives rise to a ‘funneled’ landscape that is consistent with the Lund et al study [61].

CaM’s linker flexibility appears to impact the degree to which the bottle-neck region is favored as the ionic strength is altered. In the WT linker CaM model, as the ionic strength is increased from 0.15 to 0.5 M, we see a strong redistribution of states toward the bottle-neck region. For the labile linker, this change is much less pronounced. We believe this is because the labile linker imposes less significant constraints on the CaM/CaMBR angle of approach. This is evident based on the distribution of CaMBR configurations about CaM (Fig. 3c) that fall within the bottle neck region. Notably, the CaMBR configurations are more diffusely and uniformly distributed about CaM, which suggests broader angles of approach are possible for the labile linker relative to the WT. We will further discuss how this distribution shapes association rates between CaM and CaMBR in Sect. 4.3.

We also report for WT and labile linker CaM models the number of binding events that lead to fully-bound configurations consistent with the reference crystal structure. We identify this bound state as the region near (0,0) of the native RMSD and CaMBR/CaM distance projection plane. For the WT linker CaM model, we observe ten events relative to only two for the labile linker model, out of ∼590 trials culminating in loosely-associated assemblies. This shows that a highly labile linker significantly reduces the probability of achieving the native-like bound state. It is worth speculating that this reduction in probability may have consequences in CaM’s ability to regulate its targets. This interpretation is supported by observations that CaM can bind to its targets in different structural states but only a subset of structural states can activate the target enzyme [62]..

### 4.2 Linker flexibility determines the CaM conformation ensemble

To determine the basis for how the labile linker reduces the probability of achieving native bound states, we calculated the radius of gyration (*R*_*G*_) of CaM just prior to forming the loosely-bound ensemble. In the WT linker CaM model, the *R*_*G*_ distributions can be fitted to a two-peak Gaussian distribution with maxima at 15.2 and 18.0 Å for all three ionic strengths. These two peaks correspond to a highly compact structure and an extended structure, respectively. This distribution is consistent with the extended [43] and compact CaM [63] structures that have been experimentally-observed in the absence of target. Transition path calculations have suggested that the extended CaM formation is slightly more favorable than the compact one with a ∼3-4 kcal/mol theromodynamic advantage and these two CaM states are separated by a ∼10 kcal/mol barrier [64]. Our *R*_*G*_ data of the WT linker CaM model qualitatively agree with the transition path results as the integrated extended CaM probability density is greater than that of the compact states. Moreover, the probability densities do not significantly overlap.

We contrast these data with those of the labile linker CaM model. For this configuration, a third peak emerges between the compact and extended distributions. The corresponding maxima are at 15.2, 17.0 and 18.0 Å, respectively. This additional probability density represents an intermediate state between the typical compact/extended conformation; the higher amplitude of which relative to the collapsed and extended states suggests that the intermediate state competes with two extreme CaM configurations. In other words, the alpha-helical linker of the WT model constrains the CaM conformational ensemble toward states that support productive binding. We illustrate this by labeling in green the configurations the amenable to CaMBR binding (green shaded areas in Fig. 4). It is clear from this representation that the WT linker exhibits a higher percentage of states facilitating CaMBR binding relative to the labile linker.

### 4.3 Higher linker flexibility attenuates the associate rate sensitivities to ionic strength change at elevated ionic strength

We next relate the markedly different conformation ensembles adopted by the WT and labile linkers to the kinetics of CaM/CaMBR assembly. For this, we compute a CaM/CaMBR association rate by assuming that binding is a one-step process from an unbound to a loosely-bound state. We base our assumption on observations that a fluorescence signal monitoring binding yields a monophasic distribution in time [1]. Specifically, Cook et al used an acrylodan probe that reports changes in the proteins’ hydrophobicity and polarity as they assemble. Changes in the probes fluorescence signify CaM/CaMBR association, although they do not necessarily reflect formation of the fully-bound, native-like state. Hence, we defined the CaM and CaMBR loosely-bound state by the set of configurations that adopt 50 inter-protein contacts. In Fig. 5a we verify that the loosely-bound state complexes (in gray) are thermodynamically favourable and located near the fully-bound state.

To estimate the association rate, *k*_*a*_, we report the first passage time *T*_*fp*_ of reaching the loosely-bound state in Fig. 5b,c. Our calculated *k*_*a*_ via Eq. 4 of the WT linker CaM model at 0.5 M ionic strength is 5.9 × 10^8^ M^*−*1^ s^*−*1^ (Table 1). This compares favorably to the experimental value of 2.2 × 10^8^ M^*−*1^ s^*−*1^[1] and suggests the accuracy of our strategy for computing *k*_*a*_. We demonstrate in Fig. 5c that the increased concentration of ions reduces the *k*_*a*_ values for both configurations. This is consistent with experimental measurements [1] and our Brownian dynamic simulations using isolated CaM domains and CaMBR (Fig. S1) [59]. However, we notes some differences in these trends for the WT and labile linkers. For the WT linker, increasing the ionic strength monotonically decreases the *k*_*a*_. However, this dependency is weaker for the labile linker case. We speculate that this weaker dependency is consistent with our observations that fewer labile linker configurations are confined to the bottle neck region at high ionic strengths relative to the WT (Fig. 3). This is expected as the thermodynamic stability suggested in the bottle-neck region likely contributes to the dwelling time of CaM/CaM structures in this state, which is ultimately reflected in the *T*_*fp*_. In total, our simulations indicate that the rates of forming loosely assembled CaM/CaMBR configurations are comparable for the WT and labile configurations and are strongly driven by electrostatic interactions. Importantly, though, the WT configuration strongly favors configurations that lead to fully-bound assemblies relative to the labile CaM.

**Table 1:** The mean first passage time (*T*_*fp*_) to form the loosely-bound state (Fig. 5) and the corresponding *k*_*a*_ calculated via Eq. 4

|  | WT linker |  |  | Labile linker |  |  |
| --- | --- | --- | --- | --- | --- | --- |
|  | 0 M | 0.15 M | 0.5 M | 0 M | 0.15 M | 0.5 M |
| $T_{fp}$ (ns) | 1675±156 | 2199±203 | 3442±418 | 1745±190 | 2696±301 | 2894±352 |
| $k_a$ ($10^8\text{M}^{-1}\text{s}^{-1}$) | 12.13±1.12 | 9.24±0.85 | 5.91±0.72 | 11.65±1.27 | 7.54±0.84 | 7.02±0.85 |
| Expt. $k_a$ ($10^8\text{M}^{-1}\text{s}^{-1}$) | | | 2.2 ± 0.44 | | | |

## 5. Limitations

We note a few limitations of our study that could be refined in subsequent works. In our Martini CG model setup, two assumptions were made: 1) the N-/C-domain of CaM were treated as rigid and 2) The secondary structure of CaMBR of CaN is pre-defined as an alpha-helix. The first assumption is reasonable as our comparisons of the crystal structures for CaM and CaMBR/CaM complex indicate that the structures do not significantly change (*<*2 Å RMSD). The latter assumption could mask any conformational rearrangement that the CaMBR may reflect as part of an “induced fit” binding mechanism. However, we anticipate that this limitation should only have marginal impact on our results reported here. This is motivated by the increasingly accepted notion that IDP binding mechanisms are characterized by a “couple binding and folding” paradigm comprising “conformational selection” and “induced fit” [27]. The ratio of these two mechanisms is determined by IDP’s propensity for folding in isolated state [65]. Our previous studies [59] show that when the CaMBR is simulated in the absence of CaM, it prefers partially folded *α* helix structures (68.7% of the ensemble falls within 7-9 Å RMSD of a perfect alpha-helix). We believe this implies that “conformational selection” constitutes the predominant binding mechanism for CaMBR to CaM.

In our association rate calculations, we assumed a one-step binding process based on experimental measurements. However, a two-step binding process consisting of 1) an unbound to loosely-bound (encounter complex) assembly and 2) a loosely-bound to native bound assembly through structural reorganization, is also feasible. This mechanism has been observed in other IDP binding processes [66–68]. To investigate this potential mechanism, experimental strategies that can monitor both the encounter complex and bound states are necessary. Such a strategy has been applied to melittin binding to CaM [69], for which a dansyl probe used as a fluorescence resonance energy transfer (FRET) acceptor from a melittin tryptophan residue. The approach indicated that the fluorescence signal is biphasic in time, therefore suggesting that there exists a fast rate of encounter-complex assembly, followed by a slower structural reorganization into the native complex. Analogous experimental studies for CaN binding would be helpful in elucidating this potential mechanism.

## 6 Conclusions

CaM’s flexible linker plays an important role in target binding. In this study, we utilized the Martini coarse grained (CG) model to simulate the binding process between CaM and a representative protein target, the CaMBR of CaN. Our simulations examined the fully-bound native-like CaMBR/CaM complexes under normal and high linker flexibility conditions. The latter exhibited few events that lead to fully-bound configurations, suggesting that a higher linker flexibility reducing the probability of achieving productive binding. Our data indicate that the enhanced linker flexibility disturbs CaM’s conformational ensemble in a manner that limits the sampling of states compatible with fully-bound assemblies. This limited sampling likely impacts CaM’s ability to regulate targets [62], such as the CaN phosphatase considered in our study.

Our results further highlight the importance of electrostatics interactions in driving CaMBR and CaM assembly. Notably, the nonuniform electrostatic potential along the CaM solvent exposed surface imposes constraints on angles of approach leading to protein/protein association. We demonstrate that this stereospecific constraint is more significant for the WT linker relative to the highly labile configuration, suggesting that linker lability shapes the assembly mechanism. Further, these properties influence kinetics of CaM/CaMBR assembly and their sensitivity to changes in ionic strength. In sum, our study emphasizes the important role of CaM’s linker properties in shaping the thermodyanmics and kinetics of CaM/target assembly. Our findings could therefore shed light into how CaM target regulation could be impacted by modulation of CaM’s linker properties, as might be expected for the linker-localized CaM missense mutations (M77I and S82R) [37] and post-translational modification (T80 and S82) [38].

## 7 Acknowledgements

We thank Peter Varughese for critical review of the manuscript. Research reported in this publication, release was supported by the Maximizing Investigators’ Research Award (MIRA) (R35) from the National Institute of General Medical Sciences (NIGMS) of the National Institutes of Health (NIH) under grant number R35GM124977. This work used the Extreme Science and Engineering Discovery Environment (XSEDE) [70], which is supported by the National Science Foundation under grant ACI-1548562.

## S1 Supplementary Information (SI)

**Figure S1:**
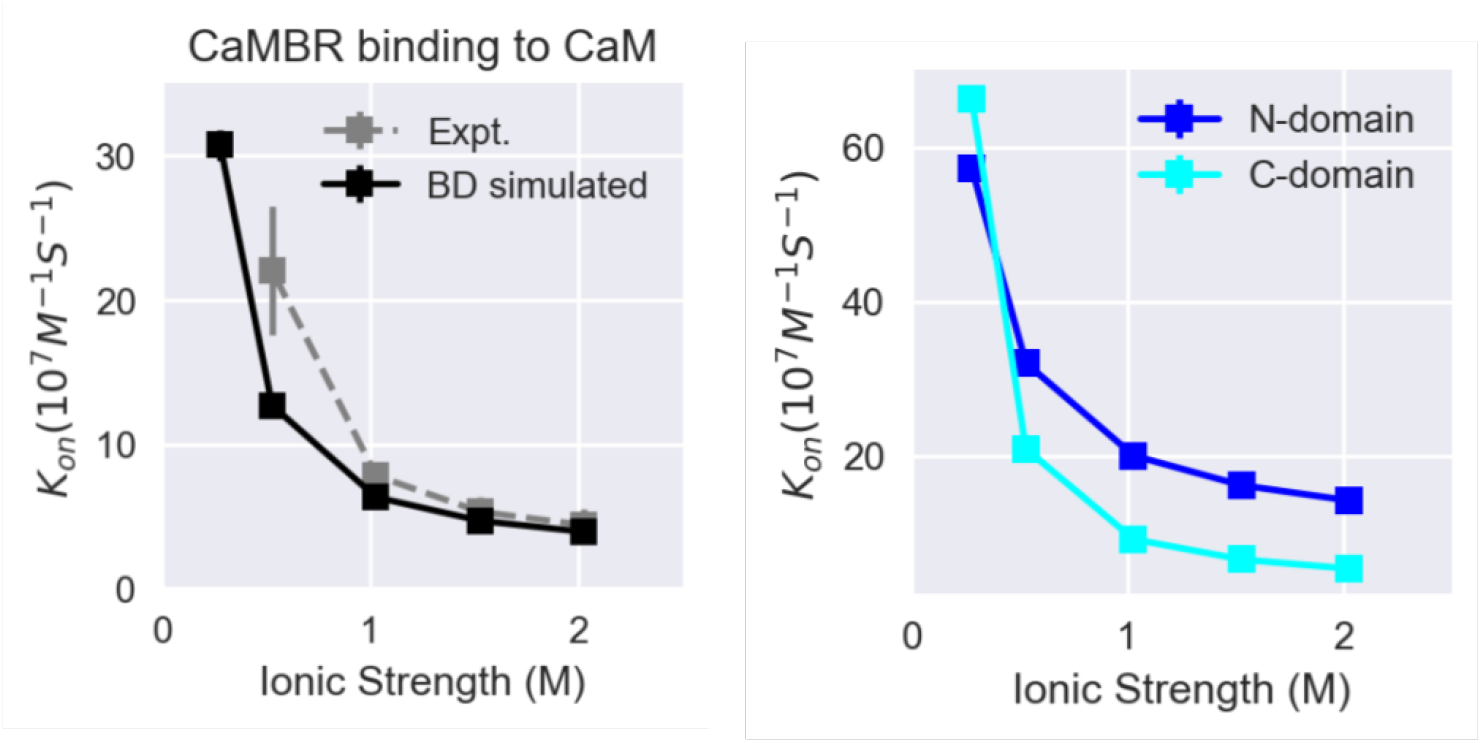
Association rate between CaM and CaMBR from Brownian dynamics (BD) simulations using the BrowDye package. In the BD simulations, the association between CaMBR and isolated CaM N-/C- domains was simulated with association rate *k*_*n*_ and *k*_*c*_, respectively. The final associate rate *k*_*a*_ is achieved via 1*/k*_*a*_ = 1*/k*_*n*_ + 1*/k*_*c*_ [59].

**Figure S2:**
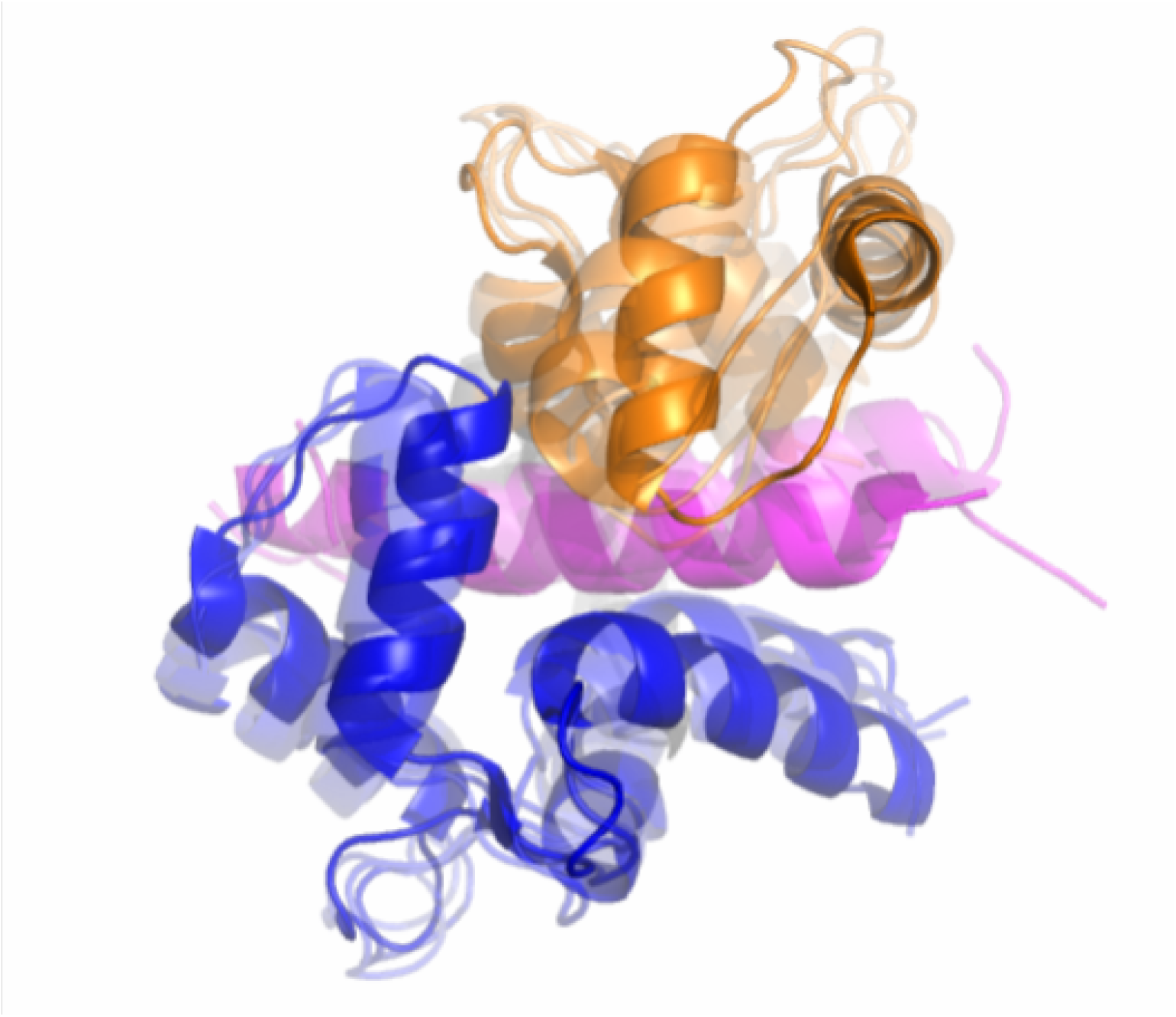
Superimposition of simulated fully-bound native-like CaM/- CaMBR complex structure with the crystal complex structure PDB 4Q5U. The simulated structures are shown as transparent cartoons..

**Figure S3:**
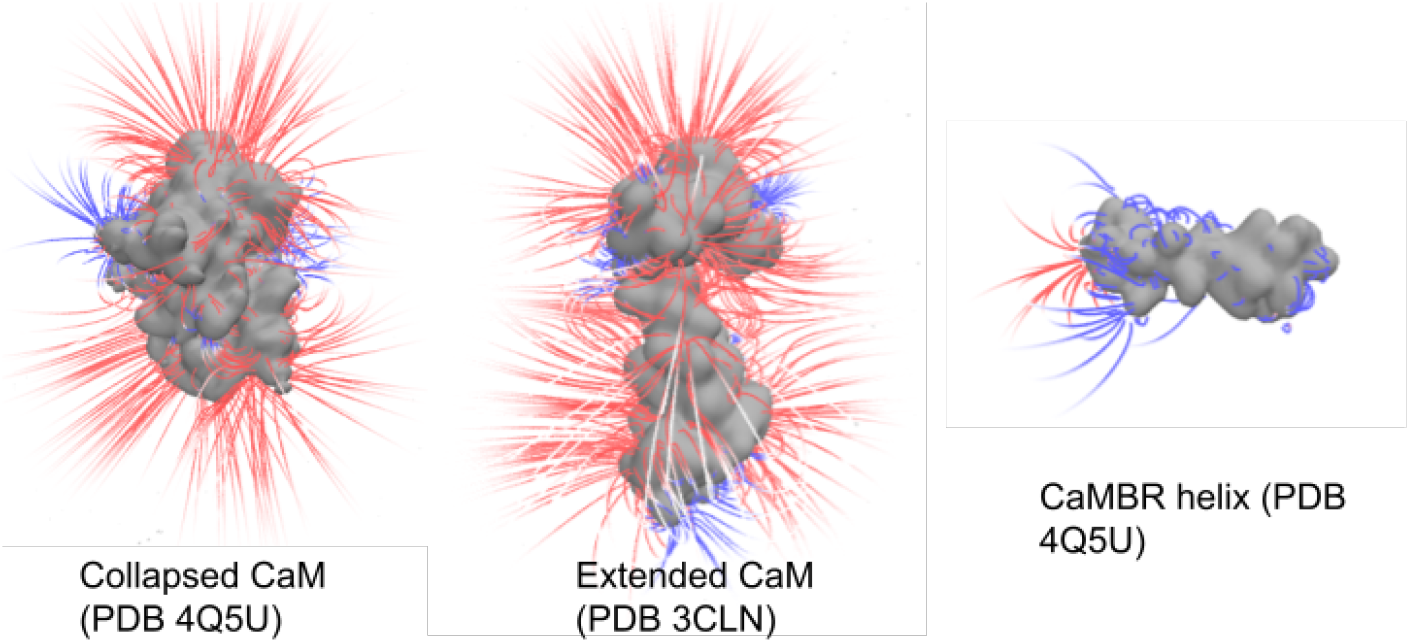
Electrostatic potentials (−0.2 to 0.2) of collapsed and extend CaM structures and the CaMBR helix. The electrostatic potentials were calculated via the APBS program at 0.15 monovalent ionic strength. The pqr files of each structure were generated using the pdb2pqr program with amber force field used.

## Notes

### Competing Interest Statement

The authors have declared no competing interest.

### Summary of Updates

The calculated ka in the abstract is mistakenly wrote as 6.3e8 M-1 s-1, the correct value is 5.9e8 M-1 s-1 from Table 1. We corrected the value in the abstract.

